# Experimental parkinsonism induced by tetanus toxin injected into basal ganglia

**DOI:** 10.1101/2024.05.09.593282

**Authors:** Patrik Meglić, Petra Šoštarić, Nikola Habek, Davor Virag, Ana Knezović, Ivica Matak

## Abstract

**Background and Objectives:** Local inhibitory circuits and long-range inhibitory projections within the interconnected basal ganglia nuclei are critical for control of voluntary movement and pathophysiology of different extrapyramidal movement disorders. Herein, we examined the major motor effects of tetanus neurotoxin (TeNT), a presynaptic neurotoxin that selectively targets the GABA-ergic synaptic transmission, when injected into individual basal ganglia nuclei.

**Methods:** The rats were injected with low-dose TeNT (0.4-0.8 ng) unilaterally into globus pallidus internus (GPi), substantia nigra (SN), or caudate putamen (CPu). The effects of TeNT-induced disinhibition were characterized by repeated assessments of motor coordination, gait, and rotational behavior, followed by measurement of regional protein content of major neuronal monoaminergic, GABA-ergic and glutamatergic population markers.

**Results:** In the beamwalk test, the CPu injection of TeNT induced contralateral plantar misplacement. TeNT injections into GPi and CPu were associated with decreased stride length and increased duration of step cycle and induced a slight ipsiversive circling during open field observation, and more intensive rotational behavior during swimming, differentially affected by D-amphetamine. Unlike rotational behavior, the gait and motor control deficits during beamwalk recovered promptly by day 14 post TeNT, which, along with the lack of reduced neuronal marker protein contents, suggested the reversibility and lack of neuronal degeneration.

**Conclusion and Implications:** Tetanus toxin injected into basal ganglia evokes transient hypokinetic motor dysfunctions consistent with experimental parkinsonism, with differential occurrence of individual motor symptoms depending on the region targeted. These results suggest that TeNT might be a useful non-neurodegenerative pharmacological agent for investigating the motor control abnormalities involving GABA-ergic basal ganglia circuits.

## 1. Introduction

Tetanus neurotoxin (TeNT) is a 150 kDa highly potent natural toxin synthesized by anaerobic bacterium *Clostridium tetani* [1] with ability to selectively affect the central inhibitory neurons and block the Ca^2+^-dependent presynaptic neurotransmitter release by proteolytic cleavage of vesicle-associated membrane protein (VAMP)/synaptobrevin [2].

The wound-associated *C. tetani* toxo-infection evokes muscle spasticity due to the neuronal entrance of a locally synthesized toxin via adjacent or distant motor nerve terminals (due to local or systemic distribution). Either way, the toxin then gets axonally transported to the spinal cord ventral horn and brainstem motor nuclei, where it selectively invades the premotor inhibitory inputs, which then causes disinhibition mediated by disbalance between excitatory (e.g. glutamatergic, cholinergic) and inhibitory transmitters (glycine and GABA) [3]. The net effect of the toxin at the level of brainstem and spinal ventral horn is uncontrollable spasm. The brainstem or spinal cord motor nuclei may not necessarily be the final destination of the toxin as it may undergo subsequent rounds of axonal transport and transcytosis within the CNS, causing disinhibition within distant interconnected regions [1,4]. With mentioned properties, the toxin itself has been employed as a neuropharmacological modulator by stereotaxic injections of the toxin into different brain regions, the most widely employed being the temporal lobe epilepsy evoked by high dose intrahippocampal TeNT [5,6,7]. While its effect in low-to-moderate *in vivo* doses is more selective for central inhibitory glycinergic and GABA-ergic terminals, at relatively high systemic or pro-convulsive doses TeNT may enter additional types of neurons, including glutamatergic and cholinergic neurons [5,8,9].

Neurological extrapyramidal disorders such as parkinsonism, Huntington’s disease and dystonias involve basal ganglia-mediated motor control impairments, associated with the disbalance of inhibitory control within and between basal ganglia nuclei. In dystonia and other hyperkinetic movement disorders, sustained or intermittent muscle hyperactivity is associated with disinhibition of thalamocortical pathway [10]. On the other hand, hypokinesia in parkinsonism and Parkinson’s disease (PD) is associated with impaired dopaminergic transmission and overt inhibition of thalamocortical pathway [10]. In the present study, we hypothesized that modulation of inhibitory control within distinct basal ganglia nuclei by TeNT may mimic some of the main motor features of mentioned motor disorders. After injection of the low non-convulsive doses of TeNT into rat unilateral GPi, SN and caudate putamen (CPu), in present study we assessed the toxin’s effects on balance, gait and rotational motor behavior, and analyzed its actions on major neuronal markers.

## 2. Methods

### 2.1. Animals

Male Wistar Han rats (University of Zagreb School of Medicine, Croatia), 6 months old, weighing 472 ± 7 g at the beginning of experiment, were used. They were kept 2–3 animals per cage in the controlled environment with a 12-hour light/dark cycle and *ad libitum* access to food and water. All procedures involving animals and animal care were conducted in accordance with European Union Directive 2010/63/EU and approved by institutional and national review boards (School of Medicine, University of Zagreb, Croatian Ministry of Agriculture, permission no. EP 229/2019). Study was reported in compliance with the ARRIVE guidelines 2.0: Updated guidelines for reporting animal research [11].

### 2.2. Pharmacological treatment

Animals were assigned randomly into six different experimental groups by block randomization, representing control vs TeNT-injected rats for each of the 3 treated regions (9 rats per TeNT injected groups, and 7 rats per corresponding vehicle injected groups). For unilateral intracerebral injection of TeNT or vehicle, the rats were deeply anesthetized with a mixture of ketamine (Ketamidor® 10%, Richter Pharma AG, Wels, Austria; 70 mg kg^-1^ intraperitoneal (i.p.) and xylazine (Xylased® Bio, Bioveta, Ivanovice na Hané, Czech Republic; 7 mg kg^-1^ i.p.) and placed in the stereotaxic apparatus with the head securely fixed. Stereotaxic delivery of 1 μL of TeNT solution (400 pg of TeNT dissolved in 1 μL of saline vehicle containing 2% BSA) via a 26G injection needle coupled to 10 μL Hamilton syringe (Cat. No. #701, Hamilton, Bonadouz, Switzerland) was performed slowly (1 μL per minute). Due to its size, the CPu-injected animals were administered two 400 pg TeNT injections into two distinct locations located rostrally and caudally within the left CPu (2 x 400 pg TeNT in 2 μl total volume). Correspondingly, sham animals were administered with an equivalent volume of vehicle. The single experimenter who performed the injection was blinded to the experimental substance administered. The stereotactic coordinates with reference to bregma were: CPu: - site 1: anteroposterior (AP) 1.3 mm, lateral (left) L 2.6 mm, dorsoventral (DV) 5.5 mm / site 2: AP -0.4 mm, L 3.6 mm, DV 5.5 mm; GPi: - AP -2.4 mm, L 2.8 mm, DV 7.8 mm; and SN - AP -5.3 mm, L 2.2 mm, DV 8.5 mm [13]. Animals were tested for potential aggravation of circling behavior by employing D-amphetamine (1 mg kg^-1^, i.p.) on day 10 after TeNT or vehicle application.

### 2.3. Behavioral tests

All behavioral tests were undertaken in a quiet laboratory room during the daytime (8-12 A.M.). In the week preceding the testing the motor performance, animals were trained to perform the beamwalk and catwalk test and establish basal performance values.

#### 2.3.1. Narrow beam walking

Rats were trained to traverse an elevated wooden narrow horizontal beam with rectangular cross-section (2.5 cm × 2.5 cm × 100 cm, elevated 50 cm above the ground), connecting a rectangular platform (10 x 10 cm) on the one side and a “safe” enclosed dark box on the other side (25 x 25 cm, with 10 x 10 cm entrance) [14–16]. The track started 10 cm after the starting platform and ending 10 cm before the “safe” box, with additional markings at 10 cm from both ends, resulting in a total of 80 cm of assessed pathway on the beam. Animals were trained one week prior to the TeNT treatment to cross the bar swiftly without tripping, halting, or losing balance. Four trials per single measurement were made, two being performed so that the camera observed the rat body right side during the passage from left to right, and the other two in the opposite direction. Trials were recorded with GoPro Hero 10 Black camera (4K, 100 fps, non-distorted linear field of view) positioned in level with the narrow beam 100 cm away from the beam to observe the entire track and entrance/exit platforms.

The analysis of video recordings enabling quantification of beamwalk transversal time and errors was done with DeepLabCut (DLC, v. 2.2.1.1) [17], an open source video tracking algorithm that allows automatic and markerless tracking of user-defined features [18], inside the Anaconda environment (Python 3.9.12). Nine distinct body parts were tracked (tail base, ear, eye, nose tip, crista iliaca, hip, lateral malleolus, metatarsus, front paw) along with two static points (START, STOP) on the horizontal narrow beam. Video pixel coordinates for the labels produced by DLC were analyzed with custom python scripts. Briefly, the time taken to cross the narrow beam was measured from when the lateral malleolus point exceeded the START point to when the front paw point surpassed the STOP point. These points were chosen to evade false start (animal hesitating before commencing the passage), as well as to exclude the brief slow down often occurring before the rat enters the closed exit platform. Slipping errors were counted only when the point overlying the metatarsus fell below the lower edge of the narrow beam (Figure 2A). Since the rat side opposite to the camera was partially obscured by the beamwalk rod, the slippage errors opposite to the camera were manually reviewed from each video.

**Figure 1.**
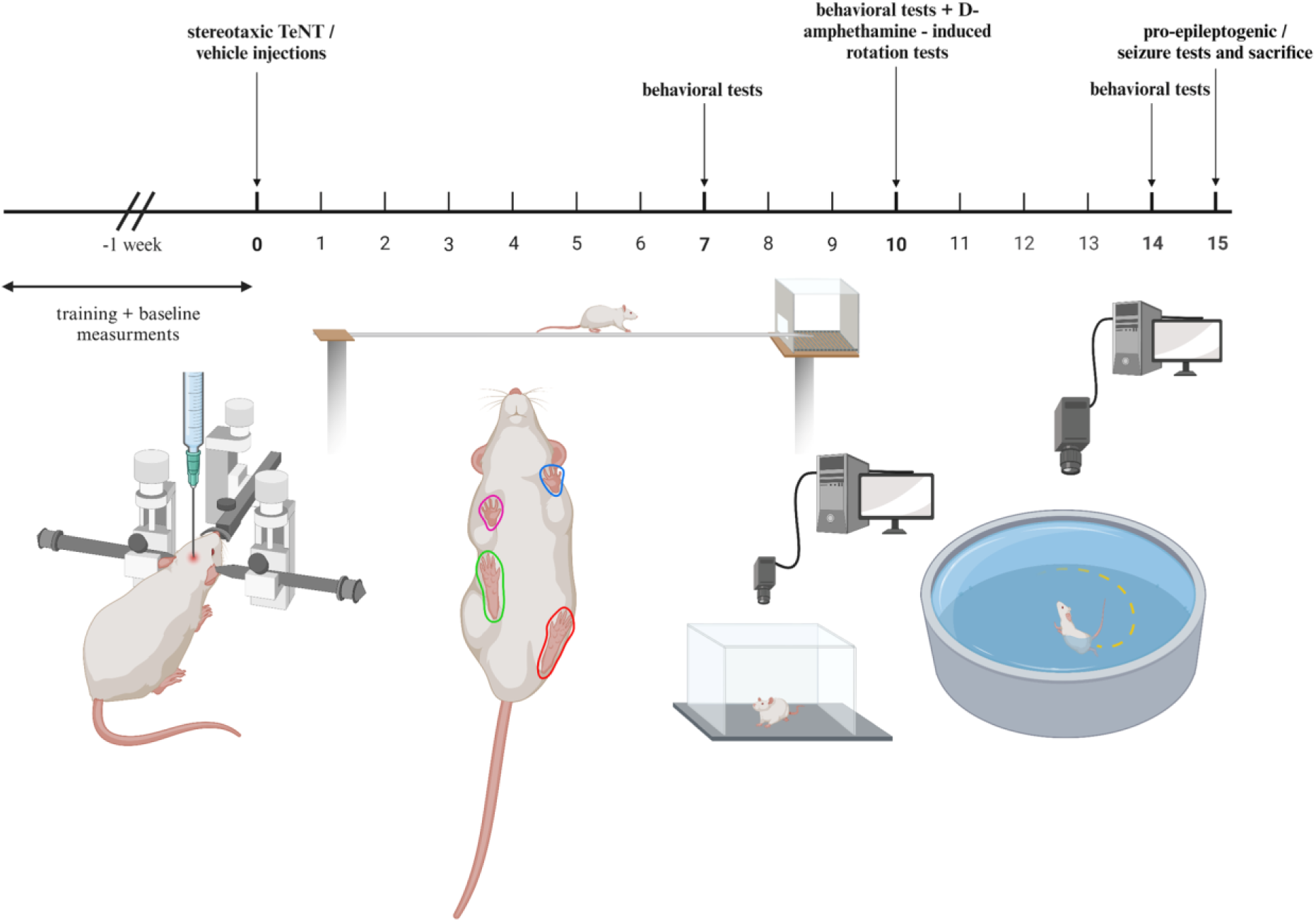
The experimental time course of treatments, behavioral assessments and schematic depiction of motor performance tests. The image was composed by Biorender (Assessed on 01/09/2024)

**Figure 2.**
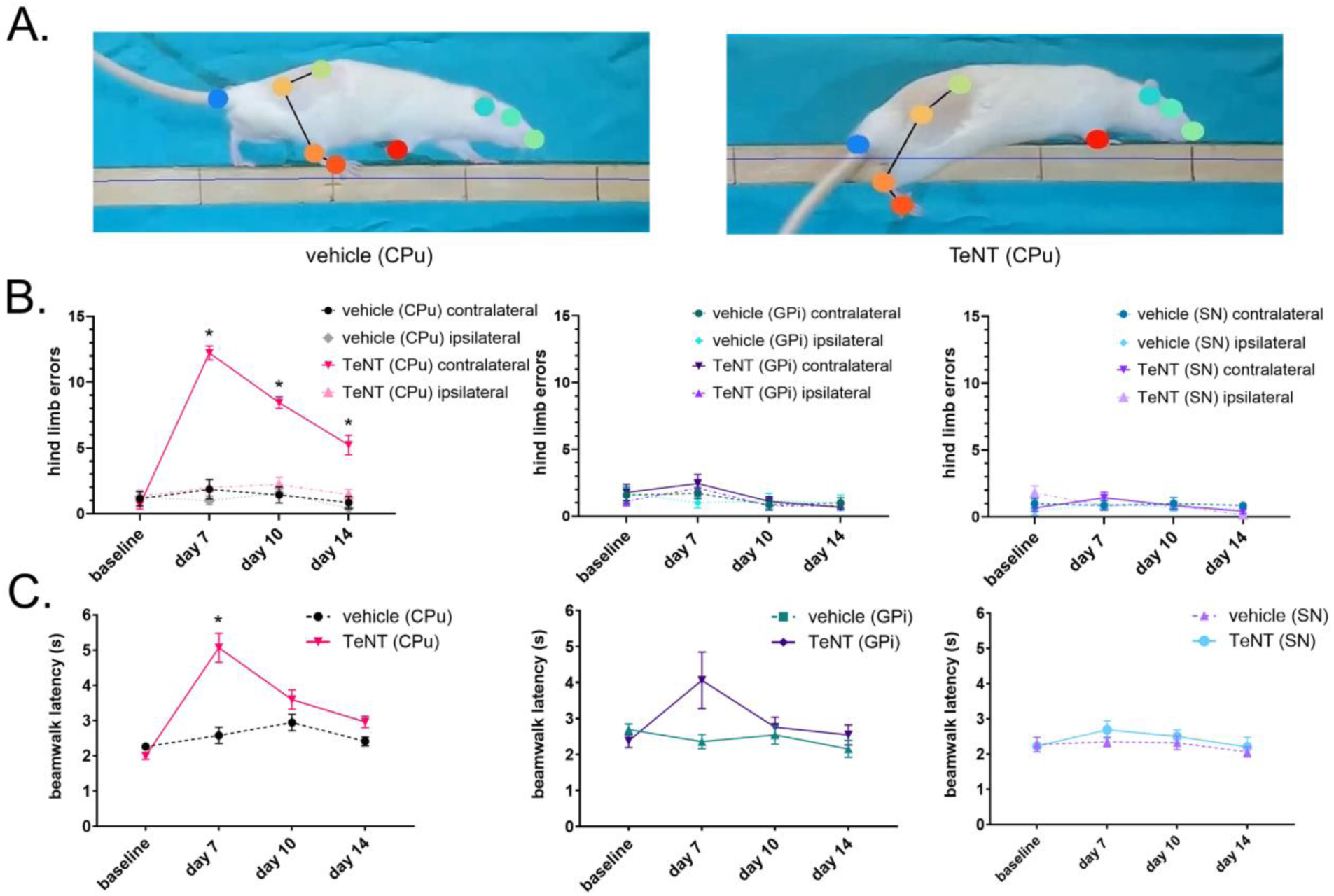
Striatal injection of tetanus toxin induces contralateral hind-limb motor control and coordination impairment during beamwalk motor performance test. Rats were injected with tetanus toxin (TeNT) into left caudate putamen (CPu, 0.8 ng TeNT), globus pallidus internus (GPi, 0.4 ng TeNT), or substantia nigra (SN, 0.4 ng TeNT). Animals were allowed to walk across the narrow beam, and the sum of hind-limb slippage errors from 4 trials, as well as the average single transition time latency, were quantified on each measurement session (baseline, days 7, 10 and 14 post TeNT). A. Markerless automatized body part recognition by DeepLabCut during normal beamwalk transversal, showing normal transition (left) and loss of balance due to right paw misplacement (right). Different body parts (tail base, ear, eye, nose tip, crista iliaca, hip, lateral malleolus, metatarsus, front paw) and the start and stop points making a line approx. 1 cm below the top surface - not visible here) were automatically labeled by different colors. B. Effect of TeNT on total number of right hind-limb (contralateral to TeNT injection) and left (ipsilateral) errors. C. Effect of TeNT on beamwalk latency. N=7-9/group; mean ± SEM; *-p < 0.05, two way RM ANOVA followed by Tukey’s post hoc.

#### 2.3.2. Open field

To assess general locomotion and rotatory behavior, animals were observed in an open field (100 x 100 cm and 50 x 50 cm). Open field box was positioned in a room with dimmed light, and the videos were acquired by GoPro Hero 10 Black camera. Rats were individually observed for 5 minutes at baseline and on 7^th^, 10^th^ and 14^th^ day after the surgery. Parameters such as distance traveled and number of rotations were quantified by using video software (Noldus Ethovision XT ver. 11.5; Noldus, Wageningen, Netherlands), The full (360 degree) body turns away from the injection side (contralateral=right i.e. clockwise) and towards the injection side (ipsilateral=left) were counted. The net rotational difference per five minutes was calculated by subtracting the total number of full anticlockwise (ipsilateral) turns from the number of clockwise turns (Δrotations=N(clockwise)-N(anticlockwise)) [19]. The D-amphetamine rotation test was performed on the 10^th^ day after the TeNT application. Animals were placed in the 50 x 50 cm transparent plexiglass box and recorded for 30 minutes in total (from 10^th^ to 40^th^ minute after i.p. application of D-amphetamine). Distance traveled and number of rotations were measured.

#### 2.3.3. Swimming

Rats were individually placed into a circular swimming pool (180 cm diameter) filled with water at depth of 30 cm maintained at 24 °C. The swimming was recorded by a wide-angle video camera (Basler AG, Basler, Ahrensburg, Germany) mounted above the pool. Video analysis software (Noldus Ethovision XT ver. 11.5; Noldus, Wageningen, Netherlands) was utilized to assess the mean and maximal swimming velocities, distance swum, mean and maximal acceleration and number of full body rotations during 120 seconds for each trial. Velocities between 10 to 100 cm s^-1^ were taken into consideration to exclude low values during passive floating or unusually high values. The parameters were calculated based on two trials per measurement. Rats were observed a few days before the toxin application and these values were considered as baseline. Same procedure was repeated on the 7^th^ and 14^th^ day after the surgery. The effect of D-amphetamine on rotational behavior during swimming was assessed on the 10th day after the TeNT application immediately after open field observation (40-50 min following the D-amphetamine i.p. challenge, two 120 second trials). Same setup and same parameters were used and measured as described previously.

#### 2.3.4. Gait analysis

The catwalk analysis was performed by employing an automated system used to assess the gait of voluntarily moving rats (GaitLab, ViewPoint, Lyon, France). The apparatus is made of a 125 cm long transparent glass runway with green LED light directed into the glass walkway sides, with the paw contact area lit up as the paw contacts the surface. This is visualized by an infrared high frame-rate video camera (100 fps). To complete each trial, the rat entered the walkway from the open end and traversed towards its home cage. A trial was considered successful if the rat completed the middle 80 cm of the walk uniformly without stopping and without any turns. At least three successful trials were recorded for each testing session. Upon training and baseline measurements, the testing was repeated on the 7^th^, 10^th^ and 14^th^ day after the surgery for all animal groups. The GaitLab software was used to analyze different dynamic (stance time, swing time, stride cycle etc.) and static gait parameters (stride length, mean intensity, paw area etc.) for each individual paw.

#### 2.3.5. Audiogenic seizures

To examine the possible occurrence of generalized epileptogenic effects of TeNT injections into the basal ganglia, a single audiogenic test attempting to provoke acoustically evoked seizures in potentially susceptible animals was performed. Animals were exposed to an acoustic stimulus (110–115 dB, 1000 Hz sound pressure levels pure tones) for a maximum of 60 seconds if no seizure occurred before [20]. The output sound signal was generated by the open source Android application Phyphox (RWTH Aachen University, Aachen, Germany) connected to loudspeakers. We observed if animals exhibited erratic running or tonic-clonic seizures.

#### 2.3.6 Prepulse inhibition of the audiogenic startle reflex

To examine possible deficits in sensorimotor gating caused by TeNT injections which might indicate hyperexcitability, a prepulse inhibition test was performed by using an open-source experimental platform (Platform for Acoustic Startle / PASTA) [21], based on conventional digital kitchen scale with its load cells connected to HX711 amplifier integrated with Arduino microcontroller board. A PC-based user interface written in Python operated via Ubuntu/Linux served as the control station which automatically plays the audio sequence containing the startle stimuli through the loudspeaker, and continuously recorded the scale measurement data at approx 80 Hz sampling rate [21]. During the testing performed in a quiet room, animals were placed in a small transparent cage covered with the wire mesh lid, secured tightly on top of the scale, and allowed to accommodate to the testing cage for 5 minutes. The test protocol involved playing a predefined sound sequence containing 10 loud white noise brief pulses (89 dB, 20 millisecond duration, separated by 10 seconds of silence between the pulses) causing startle evident as whole body linching and producing brief peaks of measured weight changes. This sequence was then followed by 10 same intensity stimuli, each preceded by a lower intensity stimulus (79 dB) 30 ms before the main pulse. The startle-evoked flinch involved rat muscle response resulting in front and hindlimb pushing against the ground, causing short peaks of increased weight measured [21]. The startle peak intensity was calculated by subtracting the rat weight from the positive peak value. Average values obtained during initial 10 startle stimuli and 10 prepulse startles that followed them were used to calculate the percentage of prepulse inhibition by using the following formula: % inhibition = (1 - mean prepulse startle (g)/mean startle (g)) x 100.

#### 2.3.7 Electroencephalogram (EEG) recording

For electroencephalogram measurement, rats (N=2) were deeply anesthetized and injected into the caudate putamen with TeNT, as explained previously. Then, small conductive steel screws (ref: 7431021, CMA Microdialysis AB, Kista, Sweden) employed as EEG electrodes were screwed into the parietal and occipital bone with the help of stereotaxic apparatus into the small craniotomy holes made by dental drill. Measurement electrode was placed over M1 motor cortex (AP -0.4 mm, L 1.75 mm), while the reference electrode was screwed over occipital bone overlying the cerebellum, behind the lambdoid suture. The two screws were soldered to the short tinned copper wire (2-3 cm) that ended on the other side with Dupont single pin female connector. After that, the scull was covered, and the electrodes connected to wire were protected by dental cement. Rats were further returned to their homecage to recover from anaesthesia and kept alone in the cage. On day 12 after TeNT application, animals were placed under isoflurane anaesthesia (5% induction). The electrodes were connected to the 36-pin wire adapter (Intan Technologies, Los Angeles, CA, USA) that was connected to the RHD 32-Channel Recording Headstage (Intan Technologies). Headstage was connected by RHD SPI Interface Cable (Intan Technologies) to the Open Ephys Acquisition Board (OpenEphys, Lisbon, Portugal). EEG signals were recorded by RHX Data Acquisition Software (Intan Technologies) with sampling rate of 20 kHz. The EEG was then recoded under various depths of anaesthesia by changing the concentration of isoflurane to read the EEG neural activity response under high level of anaesthesia (5%) for 1-2 minutes, and then the isoflurane was switched to 0% to follow the EEG neuronal activity during recovery and awakening phase (next 3-5 minutes). At least two cycles of deep anaesthesia followed by isoflurane discontinuation just before awakening were employed, and the EEG recorded. Raw data was imported in Matlab R2023b (The MathWorks, Inc., Natick, MA, US) and analysed with custom written script. Raw data were filtered using a third-order low-pass Butterworth filter to eliminate high-frequency noise. The cutoff frequency was set at 70 Hz. The filter was implemented using the bidirectional zero-phase filtering technique (filtfilt function in MATLAB), to avoid phase distortions. This method ensures the preservation of useful biological signals within the specified frequency range, while simultaneously reducing the impact of electrical noise and muscle activity artefacts. Additionally, a 50 Hz notch filter was employed to eliminate the power line interference commonly present in the recorded data.

### 2.4. Immunohistochemistry

At the end of experiment, two animals per TeNT injected group, and one animal per control group were deeply anesthetized with ketamine/xylazine and euthanized by transcardial perfusion with 400-500 ml of saline, followed by 250 ml of cold 4% buffered paraformaldehyde (PFA) fixative. Additional 2 animals were injected into the left and right GPi with 1 and 4 U of botulinum toxin type A (BoNT/A) (8 and 32 pg of 150 kDa neurotoxin) and sacrificed by perfusion/fixation after 3 days. Brains were excised and post-fixed/cryoprotected in fixative containing 15% sucrose overnight. Next day the brains were transferred to 30% sucrose in 1x phosphate-buffered saline (PBS) until they sank (approx. for additional 2 days), removed from sucrose solution and stored at -80 °C. Brains were sectioned at 30 μm thickness using cryostat (Leica) and placed in PBS-filled wells for immunohistochemistry. Free floating sections from TeNT-injected animals were quenched in PBS containing 0.25% NH4Cl for 25 minutes to reduce the endogenous tissue fluorescence, and further incubated for 2 hours in permeabilization & blocking solution (15% goat serum, 2% BSA, 0.25% gelatin, 0.20% glycine, 0.5% Triton X-100 in PBS) [22]. Subsequently, the sections were incubated with primary antibody (1:400 in blocking solution) specific for TeNT-cleaved VAMP/synaptobrevin raised in rabbit (provided by prof. Marco Pirazzini, University of Padua) [22] overnight at 4°C. After three washes in PBS, the sections were further incubated with fluorescently labeled secondary antibody (anti-rabbit IgG Fab2 Alexa Fluor 555, 1:400, 4413S/CellSignaling Technology) for 2 hours at room temperature in dark. The sections from BoNT/A-treated animals were stained by employing rabbit polyclonal antibody specific for BoNT/A-cleaved synaptosomal-associated protein 25 (anti-cSNAP-25; 1:8000, National Institute for Biological Standards and Control, Potters Bar, UK) validated in previous studies [23,24], and visualized by fluorescently labeled horseradish peroxidase substrate (tyramide-Atto 488), as previously described [16,25]. Following final washes, the sections were mounted on glass slides with anti-fading agent (Fluoromount, Sigma Aldrich) and visualized by Olympus BX-51 fluorescent microscope coupled to DP-70 digital camera and equipped with CellSens Dimension visualization and quantification software (Olympus, Tokyo, Japan). The entire coronal sections were reconstructed by employing combined multiple 4x magnification images obtained at constant exposure time. Representative images were post-processed for brightness and contrast with Adobe Photoshop.

### 2.5. Western blot

At the end of the experiment, 6-7 animals per group were deeply anesthetized with ketamine/xylazine. The animals were then decapitated and their brains removed and snap-frozen in liquid nitrogen. The brain tissue was then transferred to -80 °C for storage. Then the brains were placed inside a cooled cryostat (-25 °C). Caudate putamen was cut out manually by using the scalpel tip from thick coronal sections with the help of stereotaxic rat atlas (Paxinos and Watson, 2005) [13] as previously described [26], weighed and stored at -80 °C. Measurement of protein concentration, electrophoresis, protein visualization and transfer were done as previously described [27]. In brief, protein isolation was performed by tissue sonication inside the modified RIPA-based buffered lysis buffer, and the protein content was assessed by Lowry method. The sample proteins (20 μg) were denatured under reducing conditions, subjected to SDS-PAGE in 12% TGX stain free gel (Bio-Rad, Hercules, CA, USA), and transferred to nitrocellulose membrane by semi-dry protein transfer (TransBlot Turbo, Biorad). Following the electrophoresis and transfer, both gels and membranes were filmed by using UV gel/membrane visualization camera imager (Bio-Rad), which served as a method to visualize the total sample proteins. After the transfer, membranes were blocked for 1 h at room temperature in 5% non-fat milk in a low salt washing buffer (LSWB; 10 mM Tris, 150 mM NaCl, pH 7.5 and 0.5% Tween 20). Subsequently, they were incubated with primary antibodies, overnight at 4 °C. Primary antibodies used were anti-tyrosine hydroxylase (anti-TH; cat.#AB152, Merck Millipore, Burlington, MA, USA), anti-total-α-amino-3-hydroxy-5-methyl-4-isoxazolepropionate receptor (anti-tAMPAR) (Cat.#AB1504, Merck Millipore, Burlington, MA, USA), anti-phospho (Ser845)-AMPAR (anti-pAMPAR;cat.#AB5849, Merck Millipore, Burlington, MA, USA), anti-phosphorylated (Ser897)-N-methyl-D-aspartate receptor (anti phosphoNMDAR cat.#ABN99, Merck Millipore, Burlington, MA, USA), anti-total-calcium/calmodulin-dependent protein kinase II (anti-tCaMK; cat.#50049S, CellSignaling Technology, Danvers, MA, USA), anti-phospho-CaMK II/pCaMK (Cat.#12716S, CellSignaling Technology, Danvers, MA, USA), anti-vesicular GABA transporter (VGAT, cat.#131002, Synaptic Systems), anti-vesicular glutamate transporter 2 (anti-VGLUT2; cat.#135402, Synaptic Systems). All primary antibodies were diluted 1:1000 in the blocking solution. Membranes were washed 3 x and incubated for 1 h at room temperature with the horseradish peroxidase (HRP)-conjugated secondary antibodies (goat anti-rabbit or goat anti-mouse-HRP, 1:2000; Cell Signaling Technology, Danvers, MA, USA), washed in LSWB and incubated with chemiluminescent HRP substrate reagent (Super Signal West Femto; Thermo Fisher Scientific, Waltham, MA, USA; cat.#34095). The signals were captured and visualized using a MicroChemi video camera system (DNR Bio-Imaging Systems MicroChemi, Jerusalem, Israel). The membrane and gel protein bands were analyzed by Fiji version of ImageJ software (U.S. National Institutes of Health, Bethesda, MD, USA). The protein intensities were expressed proportionally to the signal of total proteins in the corresponding sample on the gel or membrane [27].

### 2.6. Data and Statistical Analysis

The behavioral data were represented as mean ± standard error mean (SEM). The results of repeated measurements were analyzed by two-way analysis of variance (two-way RM ANOVA) followed by Tukey’s multiple comparison post hoc test, or Sidak’s test for post hoc comparison of two groups using GraphPad 8.0 software (GraphPad Software, San Diego, CA,USA). Results of western blot data were represented by box-and-whisker plot (median, interquartile range, min/max), and analyzed by non-parametric one-way ANOVA (Kruskal Wallis test) followed by Dunn’s multiple comparison test using GraphPad 8.0. P values < 0.05 were considered statistically significant. Number of animals per group was pre-determined based on *a priori* power analysis performed with G*power software version 3.1. (University of Düsseldorf, Germany) for employed statistical test: RM ANOVA, within-between interaction; predicted size of effect: F=0.4; alpha error probability: p<0.05; power (1-β)=0.8; with estimated number of 6 per group extended to 9 due to possible attrition and need for different tissue preparation protocols (fresh tissue for western blot and perfusion-fixed for immunohistochemical analysis). The data and statistical analysis comply with the recommendations of the British Journal of Pharmacology on experimental design and analysis in pharmacology [28].

### 2.7. Materials

Following drugs were used: TeNT was purified from *C. tetani* Harvard strain cultures and was kept at -80°C [12]. When injected in vivo, the toxin was reconstituted in saline vehicle containing 2% bovine serum albumin (BSA) (Sigma Aldrich). The BoNT/A commercial preparation (INN Clostridium botulinum type A neurotoxin complex, containing 48 pg of 900 kDa neurotoxin complex equivalent to 8 pg 150 kDa neurotoxin per single mouse LD50-based international unit (1U), was dissolved in sterile saline (1 U/1 μl).

### 2.8. Nomenclature of targets and ligands

Key protein targets and ligands in this article are hyperlinked to corresponding entries in the IUPHAR/BPS Guide to PHARMACOLOGY http://www.guidetopharmacology.organd are permanently archived in the Concise Guide to PHARMACOLOGY 2021/22 [29].

## 3. Results

### 3.1. Unilateral tetanus toxins injections into the striatum induce contralateral hindlimb motor control and coordination/balance impairments

We examined the motor effect of TeNT on skilled coordinated gait during balancing by employing the beamwalk performance test. The toxin-treated animals injected into the CPu exhibited difficulties during narrow beam traversal evident as contralateral hind limb slippages (post-analyzed in slow-motion videos both automatically and by visual inspection). Apparently, the slippage error and subsequent compensatory movements to evade falling lead to the successful transversal attempts with significantly increased latency time. This impairment was particularly pronounced seven days following the TeNT injection (when almost every step resulted in slippage) and recovered incompletely until the end of the experiment, with evident improvement of paw placement resulting in lower number of errors. Despite the presence of errors, the beamwalk transversal latency recovered completely already by day 10 (Figure 2B, C). Animals injected into the GPi and SN exhibited no significant increase of errors of any individual legs, and the beamwalk traversal time was not significantly altered (Figure 2B, C).

### 3.2. Tetanus toxins injections into the basal ganglia induce transient gait impairments

Herein, we also tested the possible TeNT-evoked gait impairment by GaitLab apparatus that is able to capture multiple dynamic (e.g. step cycle & stance time duration) and static gait parameters (e.g. stride length), as well as the whole-body velocity during natural trotting gait across the narrow straight path arena. While control group animals maintained consistent velocities throughout the experiment, CPu and GPi-treated groups exhibited a significant decline in average speed on the 7^th^ day post-injections that started to recover the 10^th^ day and reached the baseline levels by 14^th^ day (Figure 3D).

**Figure 3.**
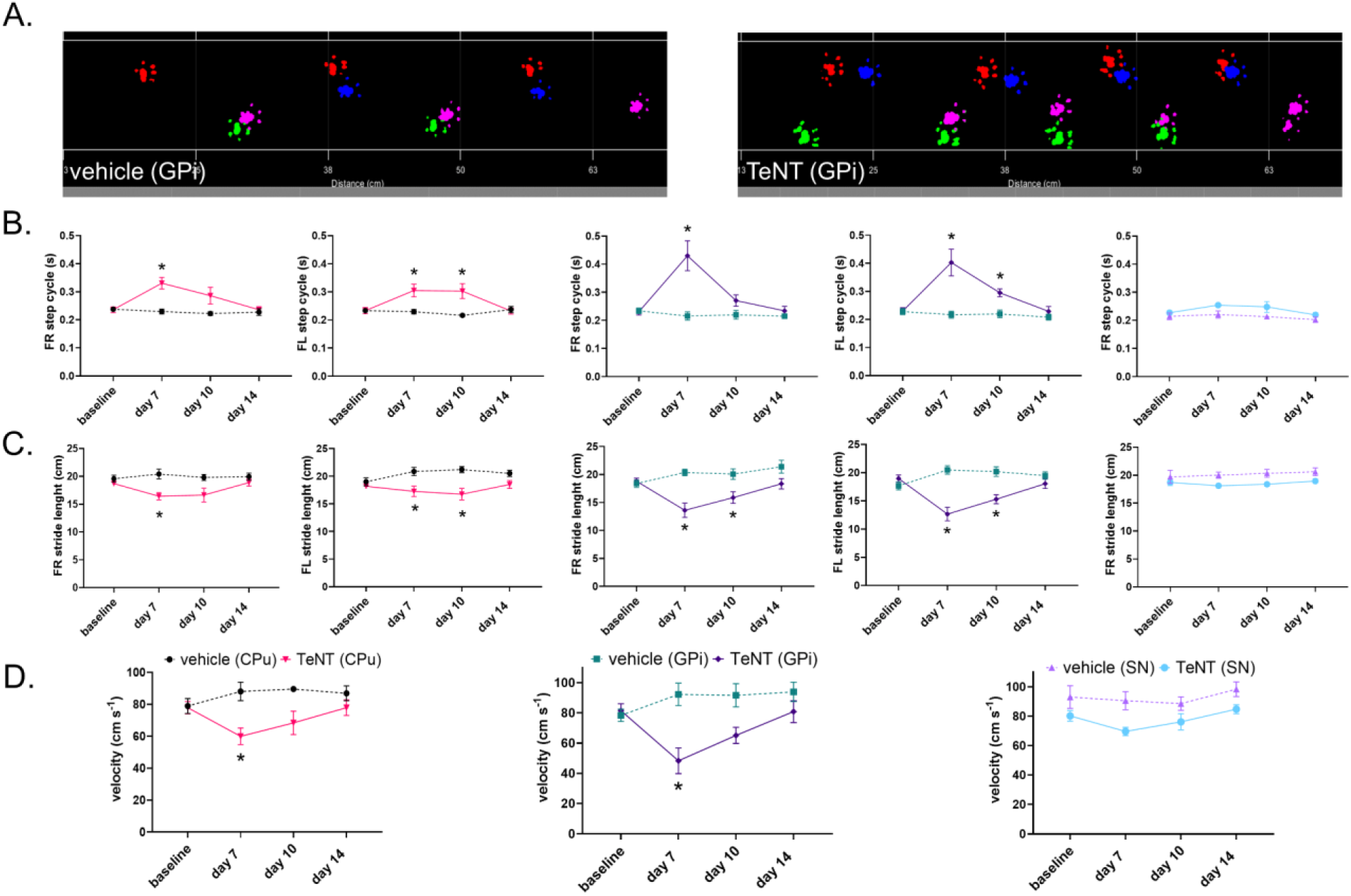
Tetanus toxin injections into the basal ganglia induce slower gait associated with transient dynamic and static gait parameter impairments. Animals were injected with tetanus toxin (TeNT) into left caudate putamen (CPu, 0.8 ng TeNT), globus pallidus internus (GPi, 0.4 ng TeNT), or substantia nigra (SN, 0.4 ng TeNT). Animals were allowed to walk across the runway, and various gait parameters from 3 trials were quantified on each measurement session (baseline, days 7, 10 and 14 post TeNT). A. Paw prints of GPi injected animal (right) and GPi control animal (left). B. Effect of TeNT on front limb step cycle (dynamic parameter). C. Effect of TeNT on front limb stride length (static parameter). D. Effect of TeNT on GaitLab-measured average whole-body velocity. FR, front right paw; FL, front left paw. N= 7-9/group; mean ± SEM; *-p < 0.05, two way RM ANOVA followed by Tukey’s post hoc.

We further looked at dynamic and static gait parameters specifically related to gait velocity, which is defined by stride length and step cycle duration roughly as a product of these two parameters. Stride length, the parameter that describes the distance between successive placements of the same paw, was reduced on the 7^th^ and 10^th^ day for all paws in CPu and GPi-injected animals (Figure 3C - only front paw values were shown since hind paw values are practically identical). In line with that, step cycle duration (corresponding to lower stepping frequency) was increased on day 7 in CPu and GPi-injected animals, and it started to recover gradually by day 14 when the values returned to basal levels (Figure 3B). Thus, the slower gait velocity was associated with reduced stride length and increased step cycle duration. No significant differences in stride length or step cycle were observed in the SN-injected group when compared to vehicle-treated controls for any paws (Figure 3B, C) on any individual day examined. However, the main column effect analysis by unpaired t-test showed significant increase in step cycle duration, decrease in stride length, and reduced average speed in TeNT vs saline-injected animals, thus suggesting a mild impairment of gait evoked by TeNT injection into the SN.

### 3.3. Unilateral tetanus toxins injections into the basal ganglia induce spontaneous ipsilateral circling behaviors

Upon animal observation and handling we observed that the body axis was tilted towards the left side, ipsilateral to the TeNT injection into CPu and GPi. The animals also evidently rotated more often towards the left side when observed in their cages with open lid, and this behavior was triggered by outer stimuli such as louder sounds or finger snapping. Thus, we analyzed more closely the animal rotation in the open field arena during normal exploration/resting, as well as during swimming inside the circular swimming pool filled with water.

Rats were recorded for 5 minutes in the open-field arena for automatic post-analysis by Noldus Ethovision. In the CPu-injected group, a slight but significant bias towards the ipsiversive rotations was observed on the day 7, 10 and 14 days after TeNT injections (Figure 4B) while for the GPi-injected animals, this bias was significant only on the 7^th^ day of the experiment (Figure 4B).

**Figure 4.**
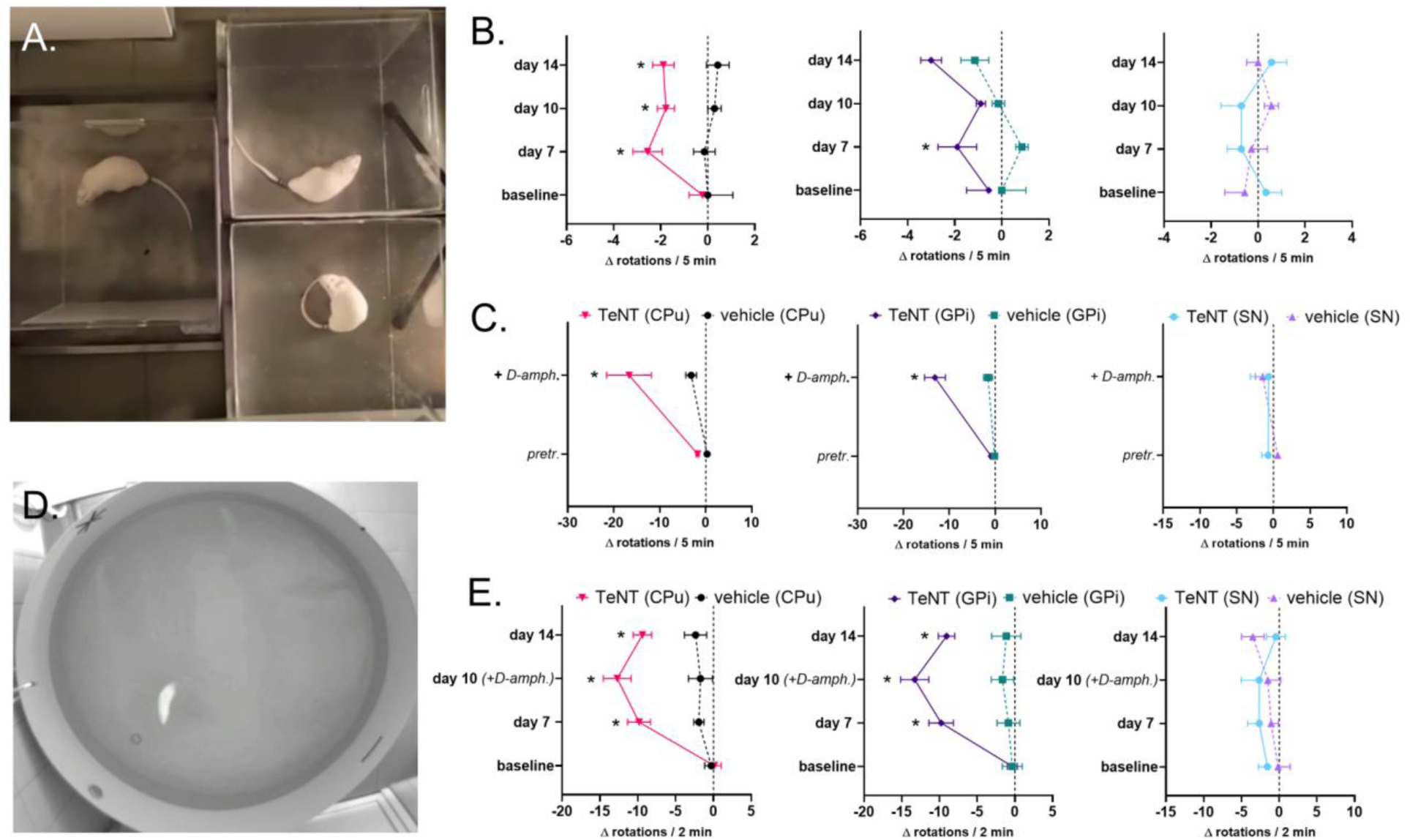
Tetanus toxin injections into the CPu and GPi induce ipsilateral rotations. Animals were injected unilaterally with tetanus toxin (TeNT) into left caudate putamen (CPu, 0.8 ng TeNT), globus pallidus internus (GPi, 0.4 ng TeNT), or substantia nigra (SN, 0.4 ng TeNT). Net rotational difference per five (open field) or two (swimming) minutes was calculated by subtracting the total number of full clockwise turns by the number of full anticlockwise (ipsilateral) turns (Δrotations=N(clockwise)-N(anticlockwise)). Rotational response to D-amphetamine was assessed 10 days after TeNT injections in the open field (10’-40’ post i.p. 1 mg kg^-1^ D-amphetamine) and upon completion of open field observation, inside a circular swimming pool filled with water (for 2 minutes). A. D-amphetamine induced rotation test in the open field. B. Effect of TeNT on spontaneous rotational behavior in the open field test. C. Effect of TeNT on spontaneous vs. D-amphetamine-induced rotational behavior in the open field test. D. Swimming test. E. Effect of TeNT on spontaneous and D-amphetamine-induced rotational behavior in the swimming test. N=7-9/group; mean ± SEM; *-p < 0.05, two way RM ANOVA followed by Tukey’s post hoc.

In the swimming test, the analysis of rotation side tendency was conducted similarly to the open field, however, with a shorter duration of test corresponding to the maximal duration of Morris Water Maze test (2 min vs 5 min open field). Compared to the open field test, the animals showed much larger spontaneous ipsiversive rotational behavior during swimming in both CPu and GPi-injected animals on the 7^th^ and 14^th^ day after TeNT injections (Figure 4E). No significant differences in rotational tendency in the open field or swimming pool were observed in the SN-injected group (Figure 4B, C).

### 3.4. D-c rotations in the open field, but not in the swimming arena

To further characterize the rotational behavior and possible involvement of dopaminergic transmission, we conducted a challenge with D-amphetamine that induces rotational behavior due to inter-cerebral hemisphere disbalance in the dopamine function in models of experimental hemiparkinsonism [19]. The rotational response to D-amphetamine was assessed 10 days after TeNT injections. On the day of the experiment, the rotations in the open field were observed before the amphetamine treatment for 5 minutes, and then starting 10 min after 1 mg kg^-1^ D-amphetamine i.p. injection, for the next 30 minutes. The analysis followed a similar approach as described for the open field and swimming tests. The final rotational difference upon amphetamine was divided by 6 to allow comparison with the 5-minute open-field tests without D-amphetamine.

The D-amphetamine administration induced several times increase in ipsilateral rotational behavior in CPu and GPi-injected animals (Figure 4C), while SN-injected group showed no significant differences (Figure 4C). Following the 30-minute open field test, the animals were placed in the swimming pool to observe the impact of D-amphetamine on rotational behavior during swimming. Although the animals were not examined before the amphetamine challenge on day 10, the values obtained with D-amphetamine were similar to the values obtained on earlier (day 7) and later occasions (day 14). Thus, it can be assumed that D-amphetamine does not augment the ipsilateral rotational behavior in the swimming test (Figure 4E). No significant changes were observed in the SN-injected group (Figure 4C, E).

### 3.5. Tetanus neurotoxin injected into the basal ganglia does not provoke pro-epileptogenic behaviors or disruption of sensorimotor gating

On the final day of the experiment (day 15 post TeNT), just before sacrifice, we examined the possible pro-epileptogenic effect of TeNT by examining the susceptibility of animals to audiogenic seizures. None of the animals exposed to an acoustic stimulus during the maximum trial duration of 60 seconds displayed any signs of wild or erratic running or epileptic seizures. In addition, we also examined the acoustic startle reflex excitability, as well as the prepulse inhibition of startle reflex. In line with the lack of anxiogenic effects of TeNT or occurrence of generalized excitability, we did not observe any differences between the startle-evoked whole body flinch intensity in TeNT-treated animals upon presentation of normal startle and prepulsed startle sequence (Figure 5).

**Figure 5.**
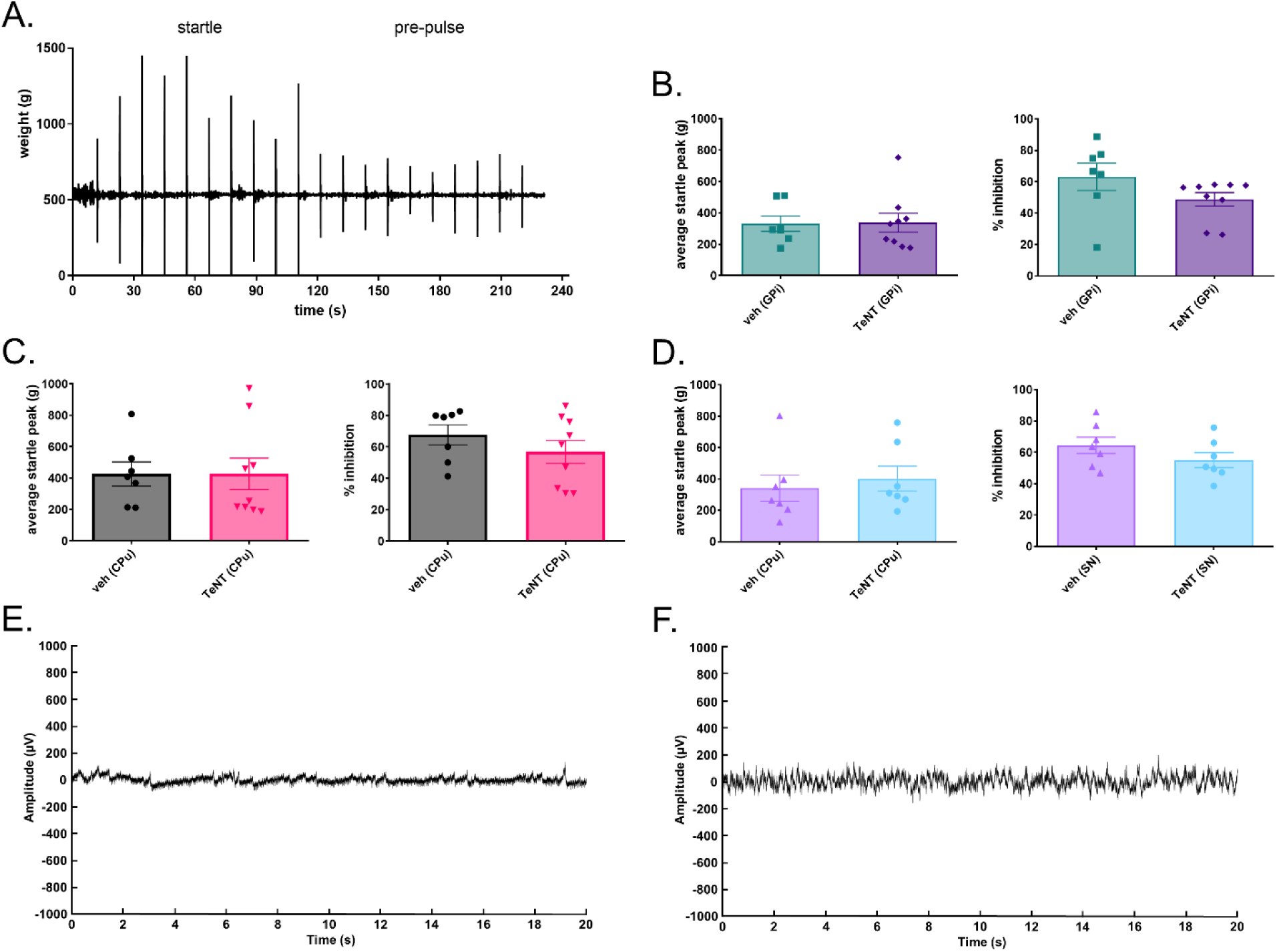
Lack of significant effect of TeNT injected into basal ganglia on audiogenic startle & prepulse inhibition, and the lack of occurrence of epileptogenic seizures. Rats were injected unilaterally with tetanus toxin (TeNT) into left caudate putamen (CPu, 0.8 ng TeNT), globus pallidus internus (GPi, 0.4 ng TeNT), or substantia nigra (SN, 0.4 ng TeNT). On day 15 post TeNT, the rats were placed in a small plastic cage secured tightly to the top of the digital scale with continuously monitored weight (80 Hz sampling rate), and then exposed to repeated white noise pulses commencing with startle-evoking sequence (approx 89 dB, 20 ms duration, 10x, 10 s delay) to establish the mean startle reflex response, and additional 10 pulses with loud pulses preceded by a short latency 10 fold lower intensity prepulse (79 dB, 30 ms prior to each loud pulse). A. Example of startle and prepulse startle responses with basal line corresponding to rat weight and peak startle calculated as positive peak deflections exerted by brief increased paw pressure on the ground by reflex flinch response. B. - D. peak startle (left panels) and prepulse inhibition responses right panels. E. EEG recording (20 s sequence) during 5% isoflurane continuous anesthesia. F. EEG recording during awakening phase from anesthesia (3-5 minutes after isoflurane concentration was set to 0% after 5%). Veh, vehicle (2% BSA in saline); % inhibition = (1 - mean prepulse startle (g)/mean startle (g)) x 100; N=7-9/group; mean ± SEM; unpaired two-tailed t-test for comparison between the two treatment groups.

On day 12 after TeNT, the rat EEG was recorded to assess for possible presence of seizure activity. In rats injected with TeNT, a normal level of EEG activity was seen, that was dependent on the depth of isoflurane anaesthesia (Figure 5E, F) without observable epileptogenic spikes. In same animals, the measurement was repeated on days 14 and 18, and no epileptogenic spiking activity was observed as well (not shown).

However, at the beginning of experiment in two animals, unilateral injection of TeNT into the SN induced intense and persistent abnormal motor behaviors evident as constant lateral barrel-rolling behaviors, starting from day 1 post TeNT. Since the animals were expected to reach the humane end-point criteria because of exhaustion and inability to feed or drink, they were immediately excluded from the experiment and sacrificed one day after the TeNT treatment.

### 3.6. TeNT evokes synaptic protein cleavage in injected and un-injected brain regions

Presently, at the end of the experiment (day 15 following TeNT) we analyzed the occurrence of VAMP/synaptobrevin cleavage at the brain coronal section level of injected sites by employing primary antibody specific for the VAMP cleavage site [22]. We found that TeNT induces robust cleaved VAMP increase on the injected side CPu (Figure 6A left panel). The part corresponding to the injection site was full of dense punctate cleaved VAMP immunoreactivity consistent with synaptic localization of the cleaved protein. In addition, punctate immunoreactivity was also observed in the form of neuronal profiles and axons in the more distant part of injected nucleus, and, occasionally in the deep layers of sensory cortex lateral to the injection site, suggesting the toxin axonal transport via projections over longer distances. No such immunoreactivity was observed in the contralateral side striatum (Figure 6 A right panel) or contralateral cortex.

**Figure 6.**
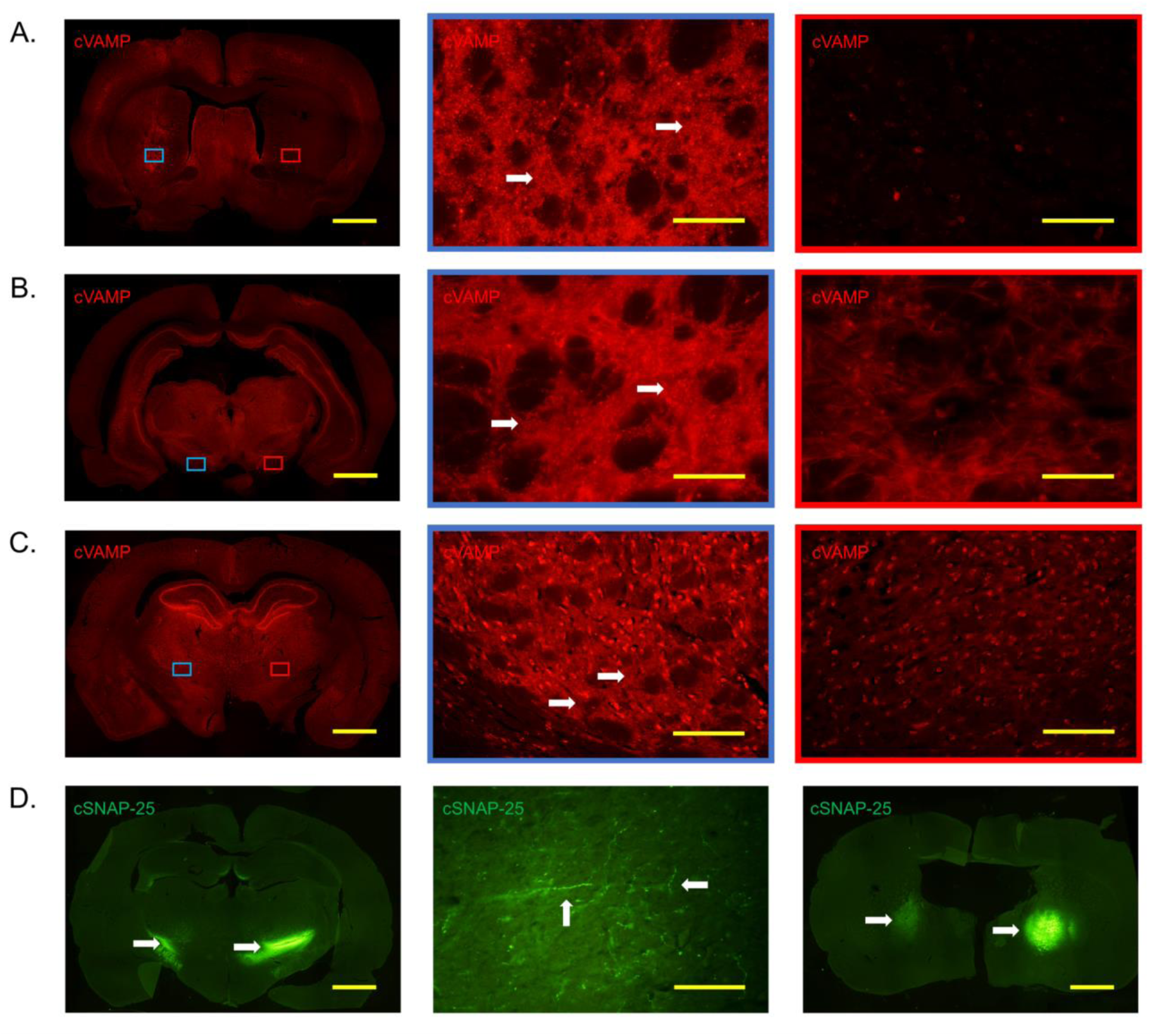
Enzymatic activity of TeNT and BoNT-A within the injected and non-injected regions. TeNT was administered into the left caudate putamen (CPu) and immunoreactivity of TeNT cleaved vesicle associated membrane protein (cVAMP) was examined on day 15 post TeNT. BoNT/A was administered into the left (8 pg of 150 kDa neurotoxin in 1 μL saline vehicle) and right (32 pg in 4 μL) globus pallidus internus (GPi), and the immunoreactivity of BoNT/A cleaved synaptosomal associated protein 25 (cSNAP-25) was examined on day 3 post BoNT/A. A. The entire coronal section showing TeNT enzymatic activity (left) in the left CPu (middle; cleaved VAMP in intensive red), and the lack of enzymatic activity in the right CPu (right). B. The entire coronal section showing TeNT enzymatic activity (left) in the left SN (middle; cleaved VAMP in intensive red), and the lack of enzymatic activity in the right SN (right). C. The entire coronal section showing TeNT enzymatic activity (left) in the left GPi (middle; cleaved VAMP in intensive red), and the lack of enzymatic activity in the right GPi (right). D. BoNT-A enzymatic activity in the left and right GPi (left), left and right globus pallidus externus (GPe) (right) and left SN ipsilateral to lower BoNT/A dose (middle). (Scale bars; A left = 1 mm, middle and right = 100 μm; B left = 1 mm, middle and right = 100 μm; C left = 1 mm, middle and right = 100 μm; D left and right = 1 mm, middle = 50 μm)

To gain more insight into the clostridial neurotoxin axonal transport following basal ganglia injection, we additionally employed injections of low dose BoNT/A, a clostridial neurotoxin homologous to TeNT and also capable of axonal transport, but forming a more stable cleavage product. Following such BoNT/A injection, we found widespread SNAP-25 cleavage in different basal ganglia regions (including globus pallidus externus, striatum and SN), thalamus and cortex (Figure 6B), thus suggesting widespread effects of clostridial neurotoxins after intracerebral injections in low-to-moderate doses.

### 3.7. The effects of TeNT on tissue protein content of neuronal monoaminergic, GABA-ergic and glutamatergic markers in striatum

At the end of experiment, we analyzed the protein tissue content of major neuronal markers: monoaminergic marker TH used to label dopaminergic nigrostriatal afferents, VGAT, an inhibitory marker that is present in inhibitory GABA-ergic terminals, and VGLUT2, marker of glutamatergic synapses. In addition, to examine possible altered activation status of excitatory glutamatergic postsynaptic markers, we analyzed the activation of ionotropic glutamate receptors by examining the ratio of phosphorylated vs total AMPAR, as well as their activator CaMKII kinase (pCaMK/CaMK), and pNMDAR. We found no reduction of either of the neuronal markers examined (Figure 7A-C), in line with lack of neurotoxic effects of TeNT. In addition, we did not observe any significant changes in the phosphorylation of glutamatergic receptor or kinase activation (Figure 7D-F).

**Figure 7.**
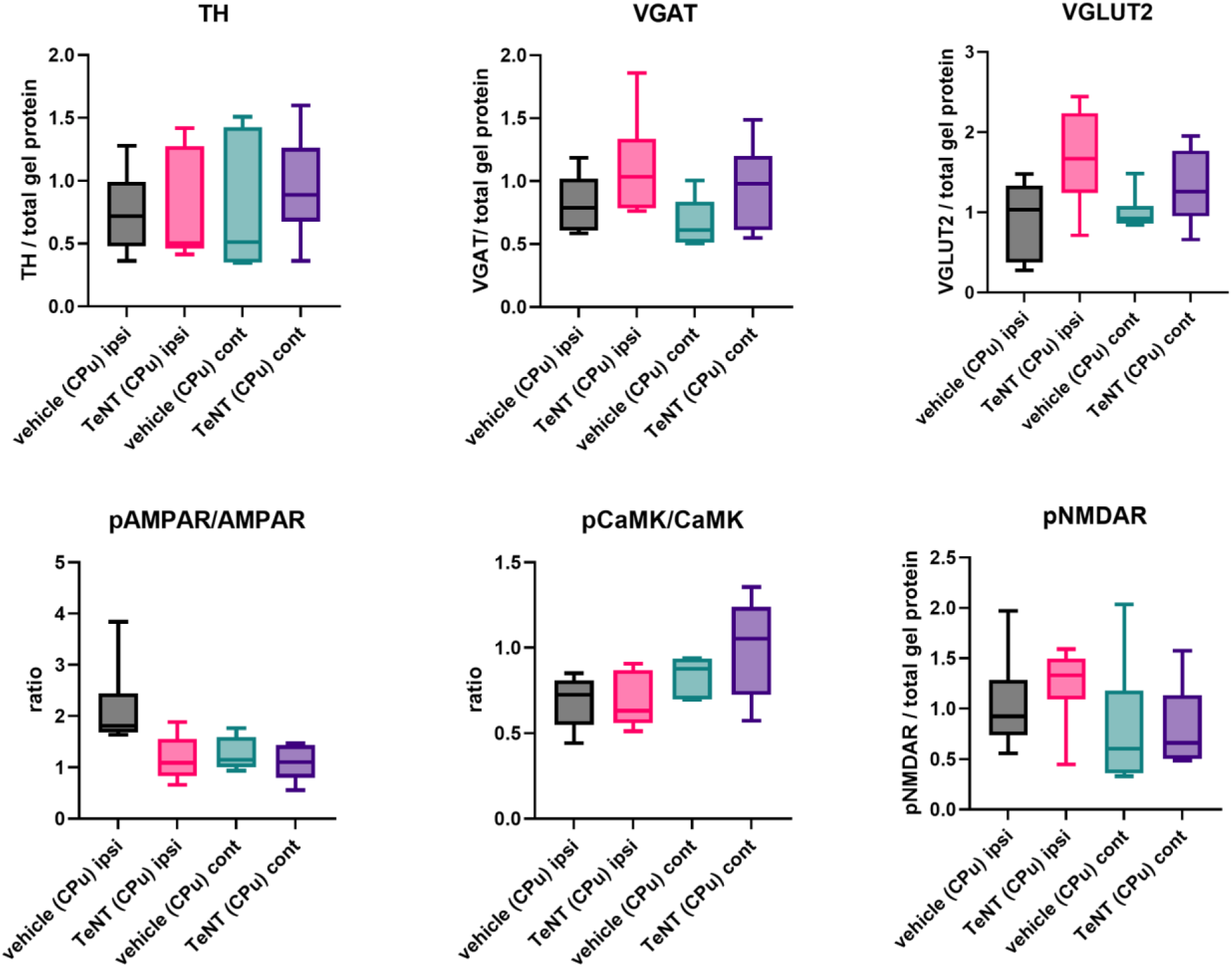
Intrastriatal TeNT does not induce significant changes in major neuronal markers and glutamatergic activation. Rats were injected unilaterally with tetanus toxin (TeNT) into left caudate putamen (CPu, 0.8 ng TeNT), and the protein tissue content of major neuronal markers and phosphorylated glutamatergic activation markers was analyzed by SDS-page and Western blot on day 15 post TeNT. A.-C. The effect of TeNT on protein markers of monoaminergic (tyrosine hydroxylase, TH), GABA-ergic (vesicular GABA transporter, VGAT) and glutamatergic neurons (vesicular glutamate transporter 2, VGLUT-2). D.-F. effect of TeNT on ratios of phosphorylated vs total AMPA receptor (pAMPAR/AMPAR), phosphorylated/total Ca-calmodulin-dependent protein kinase (pCaMK/CAMK) and the tissue content of phosphorylated NMDA receptor (pNMDAR). N = 6/group. ipsi = ipsilateral region, cont = contralateral. Data are shown as box- and whisker (median, interquartile range, min and max) and analyzed by Kruskal Wallis followed by Dunn’s post hoc.

## 4. Discussion

The basal ganglia are a set of interconnected nuclei with GABA-ergic neurons comprising the major inter-nuclear communication loops, termed direct and indirect neuronal pathways. Their main function is to regulate the inhibitory GPi/SNr output that, in turn, regulates the thalamocortical pathway-mediated initiation of the movement [30]. In parkinsonism and PD characterized by deficient nigrostriatal (SNc) dopaminergic input, the direct inhibitory pathway from striatum to GPi/SNr is suppressed, while the indirect pathway that unleashes the GPi activation by subthalamic nucleus excitatory projections is over-stimulated. Both of these effects lead to hyperactivation of GPi/SNr and, in turn, overt inhibition of the thalamic nuclei leading to hypokinesia [30]. On the other hand, the opposite role of direct and indirect pathways in dystonias and dyskinesias has been suggested to contribute to sustained or intermittent muscular hyperactivity. We hypothesized that the modulation of inhibitory control within distinct basal ganglia nuclei by TeNT may be a useful model to investigate either of these movement disorders. Although TeNT may dose-dependently target other non-GABA-ergic synapses (including dopaminergic), its effect in basal ganglia have not revealed direct presynaptic inhibition of monoaminergic transmitter release [31]. Due to its long-distance trafficking, effects of TeNT injected into basal ganglia nuclei may not necessarily be limited to the site of its injection, as it targets both the local and distant neuronal networks comprising GABA-ergic transmission. We found that TeNT, employed unilaterally in low doses into GPi and CPu, induces parkinsonism-like effects, e.g. ipsiversive circling behaviors and gait impairments (Figures 2-4).

In accordance with well-established basal ganglia connectomics, the observed parkinsonian-like motor phenotype evoked by TeNT-mediated blockage of inhibitory transmission in either CPu or GPi is in line with disinhibition of GPi and, in turn, suppressed activation of thalamic nuclei. Thus, the effect of direct GPi injection is more in line with the local disinhibitory effect of TeNT that consequently leads to increased inhibition of thalamic nuclei. While TeNT injected into CPu might enter and inhibit both indirect and/or direct medium GABA-ergic spiny neurons (iMSN/dMSN) that modulate the activity of GPi via direct or indirect pathway, the observed TeNT effects are more in line with predominating effect on dMSN and direct pathway inhibition, with consequent GPi disinhibition. Mentioned observations are in line with a recent study employing viral vector-mediated silencing of dMSN neurons in mice (by transfection of TeNT light chain) [32]. Similarly to present observations (Figure 4), in mentioned study the ipsiversive rotational behavior in the open field was exacerbated by systemic amphetamine, a sympathomimetic that reverses the transporter function and induces monoamine leakage into post-synaptic cleft. The authors argued that silencing of dMSN neurons prevents their response to dopamine-mediated stimulation arriving from SNc projections. These results are also in line with the predominating ipsi-rotational effect of dopaminergic neurotoxins such as the 6-OHDA or MPTP/MPP+ in combination with amphetamine that is related to activation of intact dopaminergic transmission within the non-treated contralateral side basal ganglia that induces motor disbalance. Thus, present observations are explainable by impaired response of GABA-ergic neurons to dopaminergic input arriving from SNc.. Interestingly, in contrast to rotational behavior in open-field, the rotational swimming behavior (that was much more pronounced compared to open-field spontaneous rotating behavior) was not significantly augmented by D-amphetamine. As a general stimulator of movement, the effect of D-amphetamine may be more exaggerated in the open field arena setting, not forcing the animals to move constantly (also performed during the inactive phase of the day when animals usually rest), in contrast to swimming task where animals need to swim constantly.

Postural instability resulting in impaired balancing and coordination leading to walking difficulties and frequent falling is recognized as one of the most incapacitating symptoms of PD. Herein we noticed that CPu-treated rats showed increased contralateral hind paw slipping errors during skilled walking across the narrow beam (Figure 2). Since quick transversal of a narrow beamwalk necessitates correct prediction of hind-limb placement during balancing, observed deficit suggests that TeNT actions in the CPu may involve deficits in limb proprioception or spatial prediction during planned movement. Interestingly, even after the hind-paw slipping, the rats did not fall from the beam by improved compensation of the transient loss of balance with body, tail and other limbs, which was evident as recovery of beamwalk latency even if hind paw slipping errors were still present. Similarly, in experimental animals, dopamine depletion leads to contralateral limb- evoked postural instability, and non-dopaminergic compensatory mechanisms [33]. Interestingly, other sites of TeNT injections (GPi and SN) did not induce contralateral hind limb errors, suggesting that such motor impairment is associated specifically with striatal circuitry.

Unlike CPu and GPi-injections, the toxin injections into SN did not induce ipsiversive turning behavior (Figure 4). In addition, the gait impairment present in CPu and GPi-injected animals were marginally altered in SN-injected animals, resulting in slight reduction of gait velocity (Figure 5). A previous study [34] showed that low dose TeNT injections induce ipsilateral turning behaviors in rats (1 -5 MLD or mouse lethal doses corresponding to 25-125 pg of presently used neurotoxin with its median lethal dose of 1 pg g^-1^). In line with mentioned earlier study, our preliminary study on few animals (n=2) injected with commercially available less potent TeNT obtained from Sigma Aldrich (Cat no. T3194) lead to ipsiversive rotational behavior. However, in the present experiment in SN-injected rats we did not observe any preference for ipsiversive vs contraversive turning. Interestingly, studies reported that GABA-ergic agonist muscimol applied into the anterior/rostral or posterior/caudal portion of SN leads to either ipsiversive or contraversive rotational behavior, and the opposite direction of turning was observed after similar injections of GABA antagonist picrotoxin [35,36]. Quickly after the injection, rostral injections of high dose TeNT may induce contraversive turning, while caudal injections may induce ipsiversive rotational behavior [37]. However, in the present more chronic study with comparably higher doses, the intranigral TeNT might have spread to affect both anterior and posterior part of the SN, which could possibly lead to cancellation of preference for ipsiversive vs contraversive rotational behavior. Compared to CPu and GPi injections, marginal alteration of gait parameters after TeNT injection into the SN could be related to the smaller effect on pathways involved in gait disturbance.

Presently, to examine the toxin’s local and distant enzymatic action in the CNS, we analyzed the clostridial neurotoxin enzymatic activity in injected and non-injected brain regions (Figure 7). In rats, the TeNT-evoked synaptic blockade is mediated by neuronal VAMP-2 isoform of VAMP/synaptobrevin, since only the VAMP-2 isoform is susceptible to TeNT-evoked cleavage [4,22]. In addition, some of the effects of TeNT in the presynaptic terminals may be associated with VAMP-2-mediated fusion events that regulate cell surface receptor exocytosis or recycling [38]. In line with its known enzymatic actions, we found the increased occurrence of cleaved VAMP-2 after TeNT injection into the basal ganglia nuclei. However, TeNT endoprotease cleaves at approximately mid-portion along the VAMP-2 molecule polypeptide [39], leading to prompt degradation of truncated synaptic protein and its rapid turnover. Therefore, a quick loss of TeNT-cleaved VAMP-2 antigen could possibly make the detection of toxin activity in uninjected distant target regions (where smaller amounts of toxin may be transported) more difficult. In comparison to truncated VAMP-2, BoNT/A-cleaved SNAP-25 is a more stable SNARE cleavage product lacking only 9 C-terminus amino acids which, in turn, can be detected more easily due to its build-up in affected neurons [40,41]. Thus, as a proof-of-principle for clostridial toxin trafficking, we examined the distribution of synaptic cleavage after low dose BoNT/A bilateral injection into the GPi (8 pg on one side and 32 pg on the other side GPi). We found that it may be transported to distant basal ganglia regions including GPe and substantia nigra, and outside of basal ganglia (thalamus, cortex). Due to its higher relative propensity for axonal transport and transcytosis in comparison to BoNT/A, it is likely that TeNT injection into GPi or CPu may lead to even higher trans-synaptic spread of the toxin to wider basal ganglia network and more regions with direct or indirect retrograde axonal connections to injected regions.

Theoretically, a possible widespread axonal and trans-synaptic spreading of the TeNT within central neurons to many different brain regions may induce elevated excitability or pro-epileptogenic/pro-seizure state. Since rodents are naturally highly sensitive to loud sounds, we examined their response to short and long-lasting audiostimuli by examining the startle response and its prepulse inhibition, as well as the possible occurrence of audiogenic seizures [42,43]. However, elevated startle response or reduced prepulse inhibition, or the occurrence of audiogenic seizures, were not observed in the experiment (Figure 5 A-D). In addition, in rats injected with TeNT into the Cpu we did not observe epileptogenic spiking activity or bursts suggestive of epileptic seizures during diffferent depths of isoflurane anesthesia (Figure 5 E, F), suggesting that the motor effects of low dose TeNT are not associated with its widespread increased CNS excitability or the occurrence of epileptic seisures.

In line with reversibility of the TeNT-evoked gait disturbances and impaired contralateral limb use/balance that quickly recover by day 14, we also did not observe any TeNT-evoked reduction of neuronal markers of glutamatergic, GABA-ergic or dopaminergic neurons in the striatum (Figure 7). Although glutamatergic transmission may be bidirectionally regulated by the GABA-ergic synaptic output [44], no changes in the phosphorylation-mediated activation of post-synaptic glutamatergic neuronal receptors and upstream kinase CaMKII were observed here at day 15 post TeNT. However, in line with the recovery of gait parameters or balance by day 14, significant effects on glutamatergic activation cannot be excluded at earlier time points. Overall, the effects of TeNT injected into basal ganglia seem to result from a non-neurodegenerative reversible synaptic blockade, making this neurotoxin a potentially useful transiently acting agent for studying the role of GABA-ergic synaptic transmission in parkinsonism-like motor dysfunctions.

## 5. Conclusion

Employed as a neuropharmacological modulator, TeNT may be used as a tool to study the function and connectivity of different motor regions in normal and impaired movement. When injected into defined basal ganglia nuclei, TeNT injections may lead to specific parkinsonian-like hypokinetic gait and balance impairments, with the extent of different motor symptoms depending on the individual nucleus targeted.

## Acknowledgments

TeNT (purified by Giampietro Schiavo and Cesare Montecucco) and primary antibody specific for TeNT-cleaved VAMP/synaptobrevin were provided by Assist. prof. Marco Pirazzini and Dr. Federico Fabris (University of Padua, Italy). Non-affinity purified rabbit polyclonal antibody to BoNT-A-cleaved SNAP-25 recognizing the SNAP-25 1–197 fragment was a kind gift from dr. Thea Sesardic and dr. Paul Stickings (National Institute for Biological Standards and Control, Potters Bar, United Kingdom).

## References

1. Fabris F, Varani S, Tonellato M, Matak I, Šoštarić P, Meglić P, et al. Facial neuromuscular junctions and brainstem nuclei are the target of tetanus neurotoxin in cephalic tetanus. JCI Insight. 2023;8:e166978.

2. Matteoli, M., Verderio, C., Rossetto, O., Iezzi, N., Coco, S., Schiavo, G., & Montecucco, C. (1996). Synaptic vesicle endocytosis mediates the entry of tetanus neurotoxin into hippocampal neurons. Proceedings of the National Academy of Sciences of the United States of America, 93(23), 13310–13315.

3. Megighian A, Pirazzini M, Fabris F, Rossetto O, Montecucco C. Tetanus and tetanus neurotoxin: From peripheral uptake to central nervous tissue targets. J Neurochem. 2021;158:1244–53.

4. González-Forero D, Morcuende S, Alvarez FJ, de la Cruz RR, Pastor AM. Transynaptic effects of tetanus neurotoxin in the oculomotor system. Brain. 2005;128:2175–88.

5. Ferecskó AS, Jiruska P, Foss L, Powell AD, Chang W-C, Sik A, et al. Structural and functional substrates of tetanus toxin in an animal model of temporal lobe epilepsy. Brain Struct Funct. 2015;220:1013–29.

6. Jefferys JG, Williams SF. Physiological and behavioural consequences of seizures induced in the rat by intrahippocampal tetanus toxin. Brain. 1987;110 ( Pt 2):517–32.

7. Mellanby J, Johansen-Berg H, Leyland R, Milward AJ. The effects of tetanus toxin-induced limbic epilepsy on the exploratory response to novelty in the rat. Epilepsia. 1999;40:1058–61.

8. Whitton PS, Britton P, Bowery NG. Tetanus toxin alters 5-hydroxytryptamine, dopamine, and their metabolites in rat hippocampus measured by in vivo microdialysis. Neurosci Lett. 1992;144:95–8.

9. Britton P, Whitton PS, Bowery NG. Effect of tetanus toxin on basal and evoked release of 5-hydroxytryptamine and dopamine in rat hippocampus in vivo. Brain Res. 1995;673:331–4.

10. Lanciego JL, Luquin N, Obeso JA. Functional Neuroanatomy of the Basal Ganglia. Cold Spring Harb Perspect Med. 2012;2:a009621.

11. Percie du Sert N, Hurst V, Ahluwalia A, Alam S, Avey MT, Baker M, et al. The ARRIVE guidelines 2.0: Updated guidelines for reporting animal research. Br J Pharmacol. 2020;177:3617–24.

12. Schiavo G, Montecucco C. Tetanus and botulism neurotoxins: isolation and assay. Methods Enzymol. 1995;248:643–52.

13. Paxinos G, Watson C. (2007). The Rat Brain in Stereotaxic Coordinates. 6th Edition, Academic Press, San Diego, USA.

14. Carter RJ, Morton J, Dunnett SB. Motor coordination and balance in rodents. Curr Protoc Neurosci. 2001;Chapter 8:Unit 8.12.

15. Matak I. Evidence for central antispastic effect of botulinum toxin type A. Br J Pharmacol. 2020;177:65–76.

16. Šoštarić P, Matić M, Nemanić D, Lučev Vasić Ž, Cifrek M, Pirazzini M, et al. Beyond neuromuscular activity: botulinum toxin type A exerts direct central action on spinal control of movement. Eur J Pharmacol. 2024;962:176242.

17. Mathis A, Mamidanna P, Cury KM, Abe T, Murthy VN, Mathis MW, et al. DeepLabCut: markerless pose estimation of user-defined body parts with deep learning. Nat Neurosci. 2018;21:1281–9.

18. Weber RZ, Mulders G, Kaiser J, Tackenberg C, Rust R. Deep learning-based behavioral profiling of rodent stroke recovery. BMC Biol. 2022;20:232.

19. Björklund A, Dunnett SB. The Amphetamine Induced Rotation Test: A Re-Assessment of Its Use as a Tool to Monitor Motor Impairment and Functional Recovery in Rodent Models of Parkinson’s Disease. J Parkinsons Dis. 2019;9:17–29.

20. Aguilar BL, Malkova L, N’Gouemo P, Forcelli PA. Genetically Epilepsy-Prone Rats Display Anxiety-Like Behaviors and Neuropsychiatric Comorbidities of Epilepsy. Front Neurol. 2018;9:476.

21. Virag D, Homolak J, Kodvanj I, Babic Perhoc A, Knezovic A, Osmanovic Barilar J, et al. Repurposing a digital kitchen scale for neuroscience research: a complete hardware and software cookbook for PASTA. Sci Rep. 2021;11:2963.

22. Fabris F, Šoštarić P, Matak I, Binz T, Toffan A, Simonato M, et al. Detection of VAMP Proteolysis by Tetanus and Botulinum Neurotoxin Type B In Vivo with a Cleavage-Specific Antibody. Int J Mol Sci. 2022;23:4355.

23. Ekong TAN, Feavers IM, Sesardic D. Recombinant SNAP-25 is an effective substrate for Clostridium botulinum type A toxin endopeptidase activity in vitro. Microbiology (Reading). 1997;143 ( Pt 10):3337–47.

24. Jones RGA, Ochiai M, Liu Y, Ekong T, Sesardic D. Development of improved SNAP25 endopeptidase immuno-assays for botulinum type A and E toxins. J Immunol Methods. 2008;329:92–101.

25. Homolak J, Babic Perhoc A, Knezovic A, Osmanovic Barilar J, Koc F, Stanton C, et al. Disbalance of the Duodenal Epithelial Cell Turnover and Apoptosis Accompanies Insensitivity of Intestinal Redox Homeostasis to Inhibition of the Brain Glucose-Dependent Insulinotropic Polypeptide Receptors in a Rat Model of Sporadic Alzheimer’s Disease. Neuroendocrinology. 2022;112:744–62.

26. Ibragić S, Matak I, Dračić A, Smajlović A, Muminović M, Proft F, et al. Effects of botulinum toxin type A facial injection on monoamines and their metabolites in sensory, limbic and motor brain regions in rats. Neurosci Lett. 2016;617:213–7.

27. Knezovic A, Piknjac M, Osmanovic Barilar J, Babic Perhoc A, Virag D, Homolak J, et al. Association of Cognitive Deficit with Glutamate and Insulin Signaling in a Rat Model of Parkinson’s Disease. Biomedicines. 2023;11:683.

28. Curtis MJ, Alexander SPH, Cirino G, George CH, Kendall DA, Insel PA, et al. Planning experiments: Updated guidance on experimental design and analysis and their reporting III. British Journal of Pharmacology. 2022;179, 3907–3913.

29. Alexander SP, Roberts RE, Broughton, BRS, Sobey CG, George CH, Stanford SC, et al. Goals and practicalities of immunoblotting and immunohistochemistry: A guide for submission to the British Journal of Pharmacology. British Journal of Pharmacology. 2018;175(3), 407–411.

30. 10. McGregor MM, Nelson AB. Circuit Mechanisms of Parkinson’s Disease. Neuron. 2019;101:1042–56.

31. Davies J, Tongroach P. Tetanus toxin and synaptic inhibition in the substantia nigra and striatum of the rat. J Physiol. 1979;290:23–36.

32. Christensen M, Nørr SE, Gether U, Rickhag M. Direct-Pathway Spiny Projection Neuron Inhibition Evokes Transient Circuit Imbalance Manifested as Rotational Behavior. Neuroscience. 2021;453:32–42.

33. Woodlee MT, Kane JR, Chang J, Cormack LK, Schallert T. Enhanced function in the good forelimb of hemi-parkinson rats: compensatory adaptation for contralateral postural instability? Exp Neurol. 2008;211:511–7.

34. Collingridge GL, Davies J. Reversible effects of low doses of tetanus toxin on synaptic inhibition in the substantia nigra and turning behaviour in the rat. Brain Res. 1980;185:455–9.

35. Reavill C, Jenner P, Leigh N, Marsden CD. Turning behaviour induced by injection of muscimol or picrotoxin into the substantia nigra demonstrates dual GABA components. Neurosci Lett. 1979;12:323–8.

36. Reavill C, Leigh N, Jenner P, Marsden CD. GABA mediated circling from substantia nigra. Nature. 1980;287:368.

37. James TA, Collingridge GL. Rapid behavioural and biochemical effects of tetanus toxin microinjected into the substantia nigra: a dual role for GABA? Neurosci Lett. 1979;11:205–8.

38. Chen H, Weinberg ZY, Kumar GA, Puthenveedu MA. Vesicle-associated membrane protein 2 is a cargo-selective v-SNARE for a subset of GPCRs. J Cell Biol. 2023;222:e202207070.

39. Schiavo G, Benfenati F, Poulain B, Rossetto O, Polverino de Laureto P, DasGupta BR, et al. Tetanus and botulinum-B neurotoxins block neurotransmitter release by proteolytic cleavage of synaptobrevin. Nature. 1992;359:832–5.

40. Schiavo G, Santucci A, Dasgupta BR, Mehta PP, Jontes J, Benfenati F, et al. Botulinum neurotoxins serotypes A and E cleave SNAP-25 at distinct COOH-terminal peptide bonds. FEBS Lett. 1993;335:99–103.

41. Bartels F, Bergel H, Bigalke H, Frevert J, Halpern J, Middlebrook J. Specific antibodies against the Zn(2+)-binding domain of clostridial neurotoxins restore exocytosis in chromaffin cells treated with tetanus or botulinum A neurotoxin. J Biol Chem. 1994;269:8122–7.

42. Curtin PCP, Preuss T. Glycine and GABAA receptors mediate tonic and phasic inhibitory processes that contribute to prepulse inhibition in the goldfish startle network. Front Neural Circuits. 2015;9:12.

43. Cunha AOS, Moradi M, de Deus JL, Ceballos CC, Benites NM, de Barcellos Filho PCG, et al. Alterations in brainstem auditory processing, the acoustic startle response and sensorimotor gating of startle in Wistar audiogenic rats (WAR), an animal model of reflex epilepsies. Brain Res. 2020;1727:146570.

44. Paraskevopoulou F, Herman MA, Rosenmund C. Glutamatergic Innervation onto Striatal Neurons Potentiates GABAergic Synaptic Output. J Neurosci. 2019;39:4448–60.

